# Is useful research data usually shared? An investigation of genome-wide association study summary statistics

**DOI:** 10.1101/622795

**Authors:** Mike A. Thelwall, Marcus Munafò, Amalia Mas Bleda, Emma Stuart, Meiko Makita, Verena Weigert, Chris Keene, Nushrat Khan, Katie Drax, Kayvan Kousha

## Abstract

Primary data collected during a research study is increasingly shared and may be re-used for new studies. To assess the extent of data sharing in favourable circumstances and whether such checks can be automated, this article investigates the summary statistics of primary human genome-wide association studies (GWAS). This type of data is highly suitable for sharing because it is a standard research output, is straightforward to use in future studies (e.g., for secondary analysis), and may be already stored in a standard format for internal sharing within multi-site research projects. Manual checks of 1799 articles from 2010 and 2017 matching a simple PubMed query for molecular epidemiology GWAS were used to identify 330 primary human GWAS papers. Of these, only 10.6% reported the location of a complete set of GWAS summary data, increasing from 4.3% in 2010 to 16.8% in 2017. Whilst information about whether data was shared was usually located clearly within a data availability statement, the exact nature of the shared data was usually unspecified. Thus, data sharing is the exception even in suitable research fields with relatively strong norms regarding data sharing. Moreover, the lack of clear data descriptions within data sharing statements greatly complicates the task of automatically characterising shared data sets.

## Introduction

Research data sharing is increasingly encouraged by funders and journals on the basis that data re-use can improve research efficiency and transparency [1,2]. For example, sharing raw data is “strongly encouraged” within open access Plan S (www.coalition-s.org). Shared data may be used to check published findings for new studies [3], for educational purposes [4] or to support further analyses [5]. The main disincentives for data sharing are a lack of access to technology or skills [6], the effort needed for curation, fears about prior publication by other researchers [7-10] and low potential for reuse in some fields [4], especially for complex non-standard datasets [11]. Nevertheless, researchers seem increasingly willing to publish their data [12]. This may generate citations to the data, originating paper or authors to recognise this effort [13-18], which is a useful incentive [19].

Not all researchers are willing to publish their data, although policy initiatives, field cultures and data infrastructure all help to encourage it [20]. The nature and extent of data sharing is highly field dependant [21]. Some fields facilitate sharing with specialised data repositories and/or standards for recording complex data (e.g., [22]). For evolutionary biology, the Dryad repository and journal data sharing mandates have combined to make data sharing almost universal in the top journals [23] (see also [24]). Such policy initiatives seem to be essential for widespread practice. Multidisciplinary generic data sharing policies from publishers can also work reasonably well, with a study of PLoS ONE finding that most articles shared data, albeit with large disciplinary differences in format and sharing method [25]. Nevertheless, most shared ecology and evolutionary research datasets are unable to be re-used due to incompleteness or practical difficulties [26]. Biodiversity datasets can be large and hybrid, combining multiple sources. The creation and citing of such datasets are supported by the Global Biodiversity Information Facility, a well-known biodiversity archive, which allows downloads of subsets of data across multiple previously published datasets and creates single DOIs to point to those subsets [27]. These may originate from unpublished work, such as routine data collection exercises or voluntary sharing [28,29], creating data quality validation concerns [30,31]. Even in fields with sharing cultures for standardised data, relatively unique datasets may not be shared, however [32]. Thus, it would be useful to know whether data sharing is widespread in conditions where it has clear value and open data publishing is supported by a research community. This is the first issue addressed in the current paper.

Data sharing may be inadequate for data re-use. The FAIR (findability, accessibility, interoperability, reusability) principles for data sharing emphasise that minimal sharing may not be effective [33]. In particular, techniques for sharing data are not widely standardised and so it is not clear whether it is possible to automatically check the extent to which data sharing occurs in any field and whether the data sharing is effective in the sense of clearly providing sufficient information for others to access, understand it. This is the second issue addressed in the current paper.

Data sharing has been increasing in biomedical science for a long time [34,35], with genomics being regarded as “a leader in the development of infrastructure, resources and policies that promote data sharing” [36] (see also [37]). Pre-publication data sharing has also been advocated in this area [38], and there are even data access committees that judge whether a team should be given access to genomic data from a controlled repository [39,40] (the “gatekeeper” model of data sharing). Within genomics, human Genome-Wide Association Studies (GWAS) seem to be particularly suitable for data sharing. These studies measure the association of genetic anomalies, generally in the form of single nucleotide polymorphisms (SNPs), across the human genome with a potentially inherited characteristic or trait of interest, such as obesity. For each individual location tested on the genome the core result is an effect size coefficient (e.g., odds ratio), standard error and corresponding *p* value derived from a test for whether a particular allele occurs more frequently among (typically) individuals in a risk group compared with a control group. The power of a test is dependent on a sample size so if two or more studies share their data and it is subsequently combined then additional SNPs may be identified [41]. Other analyses are also possible with shared summary GWAS data alone, such as cross-trait linkage disequilibrium score regression [42]. In addition, combining analyses of samples with different control groups enables more universal patterns to be discovered. This is important because many traits are influenced by multiple genes and so different sets of characteristics may produce similar outcomes in different populations. GWAS meta-analysis has evolved as a standard strategy to deal with these issues [43], although it does not seem to be widely used with shared data yet. This is slightly different from the more generic data sharing benefit of sample size increasing statistical power [44].

The GWAS data collection process is expensive and it can be time consuming due to the involvement of human subjects from which tissue samples must be taken. Thus, any data re-use has the potential to provide substantial savings in cost and time. GWAS data sharing has been mandated since January 2008 in NIH-funded research in a specific policy for this study type [45]. In practical terms, GWAS data sharing might be relatively straightforward because the key data is simple (tables of coefficients, standard errors and p values, listed against positions in the human genome using standard notation) and in large consortia data will need to be internally shared for combining, ensuring that it is typically already in a standard format. The importance of sharing GWAS summary statistics is underlined by the existence of two international databases. Whilst dbGaP (www.ncbi.nlm.nih.gov/gap) allows researchers to deposit this and related data, together with relevant metadata, the GWAS Catalog (www.ebi.ac.uk/gwas) is a manually curated record of the results of GWAS studies [46]. It includes links to public GWAS summary statistics (www.ebi.ac.uk/gwas/downloads/summary-statistic). GWAS summary statistics never contain personally identifiable information because they are cohort-wide rather than for individuals, and so they may be potentially shared publicly without privacy issues, if appropriate human subject permissions have been gained.

We assessed the prevalence of the sharing of GWAS summary statistics in published research and the potential to automatically identify this data using manual checks of 330 primary human GWAS papers from 2010 or 2017, filtered from an initial sample of 1799 papers matching a relevant PubMed query. This topic was chosen as a previously unexplored likely candidate for standardised data sharing, as well as for being a vital and vibrant research area. The years 2010 and 2017 were chosen to help reveal changes over time. The following research questions encapsulate the broad goals of the project.

1. What proportion of primary human GWAS share data?
2. Can shared primary GWAS data be automatically identified?

## Methods

A PubMed query was used to identify articles likely to be primary GWAS. PubMed was used since its scope should encompass most GWAS journal articles. A simple query was used rather than a more complex version to enable easier interpretation of the results of the article identification stage. The query was as follows, where the term molecular epidemiology was added to filter out methods-based articles.

**“Molecular Epidemiology”[Majr] AND “Genome-Wide Association Study”[Majr]**

After discarding papers that had types other than research-article, this gave 867 journal articles from 2010 and 932 from 2017. The year 2010 was chosen as the first year with close to the maximum number of GWAS per year. The year 2017 was selected instead of 2018 (in January 2019, at the time of data collection) because there were fewer articles in 2018 than in 2017, suggesting that some PubMed records from this year were missing. The articles were checked for being primary human GWAS by three experienced content analysts by reading their titles, abstract or full text until the classification was clear. The process was as follows.

Articles for non-human genomes were discarded. Articles with the term “meta-analysis” were initially all classed as primary GWAS and then checked by a GWAS expert (MM). In contrast to general meta-analyses, GWAS meta-analyses are usually primary studies that analyse, at least in part, freshly collected data from multiple cohorts. Here, “meta-analysis” refers to the combination of data from multiple sources (i.e., different study samples) rather than a secondary analysis combining data from previously published sources. Articles that were difficult to categorise were forwarded for checking by a GWAS expert (MM). It was not straightforward to check whether an article reported a primary GWAS because it may include both primary and secondary GWAS, it may include prior, parallel or follow-up experiments or analyses, and the details may be described in technical language that avoids the term GWAS within the methods and results. The first author re-checked all articles classified as primary human GWAS. A fifth coder, a GWAS researcher (MM) checked 77 random articles and made 12 corrections (16%), in all cases ruling out an article initially judged to be a primary human GWAS. For example, “Locus category based analysis of a large genome-wide association study of rheumatoid arthritis” had been categorised as a primary human GWAS because the initial coder and follow-up check had not detected that it did not report primary data.

After identifying an article as primary human GWAS, the same set of three coders attempted to identify whether it shared GWAS summary statistics. This information was first sought in Data Availability statements, if any, or at the end of the article, or in associated supplementary information files. Failing these, the remainder of the article was scanned for references to summary statistics. An article was recorded as sharing summary statistics only if it included a complete set rather than just the statistically significant ones because a full set is needed for re-use. A fourth coder (the first author) checked these results and extracted the text in each article referring to the summary data.

As an additional follow-up check, articles in the European Bioinformatics Institute GWAS Catalog (www.ebi.ac.uk/gwas) from 2010 and 2017 with public summary statistics were cross-referenced with the main data set investigated and reasons for any differences identified. This revealed some mismatches and one clear mistake. The original search had missed some primary GWAS without MeSH terms and that had not been matched to *Molecular epidemiology* by PubMed. One matching article had been classed as non-primary GWAS in the main dataset thorough human error.

All identified GWAS Summary Statistics were examined for associated metadata. Without effective descriptions, data is harder to reuse [47].

## Results

A minority (18.3%) of the articles matching the MeSH query were judged to be primary human GWAS. Other articles matching the query were about animals or plants (n=252), or were commentaries, methods-based, used different methods (e.g., whole genome sequencing), or were follow-up studies. Often the GWAS status of an article was not clear from its title and methods details had to be checked. For example, “Genetic association study of exfoliation syndrome identifies a protective rare variant at LOXL1 and five new susceptibility loci” was classified as non-GWAS because it was not genome-wide, “we collected a global sample of XFS cases to refine the association at LOXL1”. The articles classified as primary human GWAS were analysed for the presence of information about the availability of GWAS summary statistics.

### Availability of GWAS summary statistics

Out of all 330 articles classified as primary human GWAS, 10.6% reported sharing GWAS summary statistics in some form, increasing substantially from 4.3% in 2010 to 16.8% in 2017 (Table 1). If an article did not state that its data was shared, it may still be possible to email the authors to access it or the authors may have subsequently deposited it elsewhere after publication. Conversely, data sharing promised by the authors may not materialise in practice (and perhaps rarely does: [48]) and is time limited. In addition, data sharing statements often did not specify the type of data, so those that were offered by email or by request may not include complete GWAS summary statistics.

**Table 1.** Availability of summary statistics in published primary GWAS articles from 2010 and 2017, according to the article text.

| **GWAS summary statistics availability** | **2010** | **2017** | **Total** | **Percent** |
| --- | --- | --- | --- | --- |
| Not stated in article | 156 | 139 | 294 | 89.4% |
| Broken link or not findable at stated location | 3 | 1 | 4 | 1.2% |
| On request to the authors | 0 | 8 | 9 | 2.4% |
| On request via dbGaP | 2 | 5 | 7 | 2.1% |
| On request via EGA | 1 | 3 | 4 | 1.2% |
| On request via another portal | 0 | 3 | 3 | 0.9% |
| Free online without login, proprietary format | 1 | 0 | 1 | 0.3% |
| Free online without login, plain text | 0 | 8 | 8 | 2.4% |
| **Human GWAS with primary GWAS data** | **163** | **167** | **330** | **100.0%** |
| Articles checked | 867 | 932 | 1799 |  |

When data sharing was flagged in an article, a variety of strategies could be used. Most data required a permission seeking stage, either directly from the authors (2.4%) or through the database of Genotypes and Phenotypes (dbGaP) or the European Genome-phenome Archive (EGA) or another access-controlled portal, all of which have approval processes that must be completed before the data can be accessed (4.2%). The summary statistics were open access in a non-proprietary format in only 2.4% of cases, with 2 of these 8 cases lacking descriptive metadata. Thus, when shared, some form of data access control is usually employed.

### Descriptions of the availability of GWAS summary statistics

Articles sharing GWAS summary statistics usually reported this in a Data Availability section within the article (26 out of 35), although two were mentioned in other relevant sections (Author Information, Supplementary Information). In these sections the location of information about shared data should be straightforward to identify because the sections are short and focused on this goal. Such information would be more difficult to automatically extract when mentioned in the methods (4 articles) and results (3 articles) sections because it would first need to be identified and then delimited from the rest of the hosting section.

Only five data sharing statements directly described the shared data as GWAS Summary Statistics (bold and italic in the list below), and all five used different phrases. The more general term “genotype data” (found in 8 articles) was more common. This term is ambiguous because there are other forms of genetic analysis. Just under half of the articles describe the sharing policy in the most indirect manner possible, with anaphors “datasets” or “data” (used in 16 articles). Since most articles typically employed multiple analyses and might share incomplete datasets (e.g., just the top SNPs identified, or with the results from some study samples removed), a dataset would need to be identified, downloaded and inspected to check whether it contained complete GWAS summary statistics. In some cases, the data sharing link was to a project website containing similar data from multiple studies so article title matching in the target site was needed to identify the correct dataset. Thus, it would be difficult to automatically identify from data sharing statements whether GWAS summary statistics were shared. The following is a complete list of data sharing statements, together with an indication of where they occurred in each article.

1. **Authors:** “Data Availability: [] **Data** requests can be made by contacting []”
2. **Authors:** “Data availability: Due to data protection issues, the **raw data** cannot be made publiclly available. However, individual researchers may request to use the data for specific projects on a collaborative basis.”
3. **Authors**: “Data availability: Genotype data of GERA participants are available from the dbGaP (database of Genotypes and Phenotypes) under accession phs000674.v2.p2. This includes individuals who consented to having their data shared with dbGaP. The complete GERA data are available upon application to the KP Research Bank (https://researchbank.kaiserpermanente.org/). The **summary statistics** generated in this study are available from the corresponding authors upon reasonable request. The GWAS summary statistics for the replication study11 are available from (https://www.dropbox.com/sh/3j2h9qdbzjwvaj1/AABFD1eyNetiF63I5bQooYura?dl¼0).”
4. **Authors:** “Data availability: The ***full GWAS summary statistics*** for the 23andMe discovery data set may be requested from 23andMe, Inc. and received subject to the execution of 23andMe’s standard data transfer agreement, which includes clauses intended to protect the privacy of 23andMe participants, among other matters.”
5. **Authors (also dbGaP and EGA):** “Data availability: The scan IARC-2 obtained Institutional Review Board certification permitting data sharing in accordance with the US NIH Policy for Sharing of Data Obtained in NIH Supported or Conducted GWAS. Data are accessible on dbGaP (study name: ‘Pooled Genome-Wide Analysis of Kidney Cancer Risk (KIDRISK)’; url: http://www.ncbi.nlm.nih.gov/projects/gap/cgi-bin/study.cgi?study_id=phs001271.v1.p1). Similarly, the NCI-1 scan is accessible on dbGaP (phs000351.v1.p1). Data from IARC-1 and MDA scans are available from Paul Brennan and Xifeng Wu, respectively, upon reasonable request. The UK **scan data** will be made available on the European Genome-phenome Archive database (accession number: EGAS00001002336). The NCI-2 scan will be posted on dbGaP.”
6. **Authors:** “Data Availability Statement: Please contact author for **data** requests”
7. **Authors:** “Data Availability Statement: The Icelandic population [Whole Genome Sequencing] data have been deposited at the European Variant Archive under accession code PRJEB8636. The authors declare that the **data** supporting the findings of this study are available within the article, its Supplementary Information files and on request.”
8. **Authors:** “Availability of data and materials: The **dataset** generated in AA-DHS are available from the senior author based on reasonable request. JHS [replication] data are available on dbGap and/or direct request addressed to the JHS leadership.”
9. **dbGaP**: “Data Availability: [] The [Health and Retirement Study] **genotype data** is available to approved users through the NCBI Database of Genotypes and Phenotypes (dbGaP).”
10. **dbGaP**: [in Methods section] “The **datasets** used for the analyses described in this paper can be obtained from the database of Genotypes and Phenotypes (dbGaP) at http://www.ncbi.nlm.nih.gov/projects/gap/cgi-bin/study.cgi?study_id=phs000092.v1.p1 through dbGaP accession number phs000092.v1.p1.”
11. **dbGaP**: “Data Availability Statement: **Genotype data** from the GICC GWAS are available from the database of Genotypes and Phenotypes (dbGaP) under accession phs001319.v1.p1.”
12. **dbGaP:** [in Methods section] “**Meta-analysis results** are available on dbGaP (https://www.ncbi.nlm.nih.gov/gap; accession number phs000930).”
13. **dbGaP:** “Data Availability: The primary **data** are available from dbGAP, accession number phs000431.v2.p1.”
14. **dbGaP:** “Data availability: We have deposited all **genotype data** supporting our findings from the discovery cohort in the Database of Genotypes and Phenotypes (dbGaP), with accession code phs000421.v1.p1. Other data that support our findings are available from the authors by request; see author contributions and their published references for specific data sets.”
15. **dbGaP**: “Data accession: The **genotype data** for the 311,459 SNPs in 1215 Behçet’s disease cases and 1278 healthy controls from Turkey have been deposited in the National Institutes of Health database of genes and phenotypes, dbGaP (http://www.ncbi.nlm.nih.gov/sites/entrez?db=gap), accession number: phs000272.v1.p1”
16. **EGA:** “Data Availability Statement: A list of the SNPs in the discovery scan exhibiting P < 10-4 are available in Supplementary Table 9. Researchers can gain access to the **data** by applying to the data access committee (www.ebi.ac.uk/ega/).”
17. **EGA:** “DATA AVAILABILITY: ***Case Oncoarray GWAS data*** and the Hi-C dataset utilized in this paper have both been deposited in the European Genome–phenome Archive (EGA), which is hosted by the European Bioinformatics Institute (EBI), under the accession codes EGAS00001001836 and EGAS00001001930 respectively.”
18. **EGA:** “Data Availability: All relevant **data** are available from the European Genome Archive (EGA) with the accession number: EGAD00010001447.”
19. **EGA**: “Availability of data and materials: All **data** sets generated as part of this study are available at the European Genome-phenome Archive (EGA) [89] under the following accession numbers: EGAS00001001456 for 450 K array data.”
20. **Other portal**: “Data availability: Data, including all **genotype data** and information on hypertension status, are available on approximately 78% of [Genetic Epidemiology Research on Adult Health and Aging] participants from dbGaP under accession code phs000674.v1.p1. This includes individuals who consented to having their data shared with dbGaP. The complete GERA data are available upon application to the KP Research Bank Portal”
21. **Other portal**: “Data availability: Stage one **data** are from UK Biobank, and can be obtained upon application (ukbiobank.ac.uk)”
22. **Other portal:** [in Materials and Methods section] “The **raw genotype and phenotype data** of the Tibetan and Han subjects are available through application at https://www.wmubiobank.org”.
23. **Open access proprietary:** [in Methods section] “GTYPE was used for analysis of signal intensity and for genotype calling (Affymetrix; **full SNP genotype data** are available at www.icr.ac.uk/array/array.html)”
24. **Open access**: “Data Availability: A **dataset** file is available from the GRASP resources data. The URL is https://grasp.nhlbi.nih.gov/FullResults.aspx. The study will be found using the first author name (Salem) or the pubmed ID.”
25. **Open access:** “Data Availability: All relevant **data** are within the manuscript, supporting information files, and hosted at the following URL: http://cmgm.stanford.edu/~kimlab/ACL/Achilles_ACL.html. Data will also be available at NIH GRASP: https://grasp.nhlbi.nih.gov/FullResults.aspx.”
26. **Open access:** “Data Availability: All relevant **data** can be accessed at NIH GRASP by using the following link: https://grasp.nhlbi.nih.gov/FullResults.aspx.”
27. **Open access**: “Data Availability: All the **summary level data**, as well as the individual level data for Tanzania are available from DRYAD (doi:10.5061/dryad.cq183)”.
28. **Open access:** “Data Availability: These third party **data** are available from NIH GRASP. The authors did not have any special access privileges and interested researchers can access the data at https://grasp.nhlbi.nih.gov/FullResults.aspx (Trait(s): Ankle injury).”
29. **Open access**: “Data Availability Statement. ***Summary GWAS estimates*** for the T2D meta-analysis and bivariate summary data are publicly available at the following:”
30. **Open access:** “Data availability. The **genotype data**, BMI measurements, and related phenotype information that support the findings of this study are available in Japanese Genotype-phenotype Archive (JGA) under accession codes JGAS00000000114 for the study, JGAD00000000123 for the genotype data, and JGAD00000000124 for the BMI measurements. The summary statistics of the GWAS have been deposited in the National Bioscience Database Center under data set identifier hum0014.v6.158k.v1.”
31. **Open access:** “Supplementary information: [] Supplementary Data 2: ***Summary statistics for the genome-wide association study***.”
32. **Missing/broken**: [in Results section] “The complete set of **results from this genomewide association study** can be found in the National Institutes of Health Genotype and Phenotype database (dbGaP; www.ncbi.nlm.nih.gov/projects/gap/cgi-bin/about.html) (accession number phs000233.v1.p1).”
33. **Missing/broken**: [in Results section] “A ***genome-wide set of summary association statistics*** will be available at the National Bioscience Database Center (NBDC)”
34. **Missing/broken**: [in Results section] “The **summary of statistical analysis in the first stage** is available on the genome-wide association database (https://gwas.lifesciencedb.jp/cgi-bin/gwasdb/gwas_study.cgi?id=cerebral).”
35. **Missing/broken**: “Author Information: [] Full **data** are available, under a data access mechanism, from the European Genome-phenome Archive (http://www.ebi.ac.uk/ega/page.php).”

## Limitations

The results are limited by the initial MeSH query used, which did not match all GWAS studies, and the use of non-expert coders and cross-checker to classify most of the articles. The results are also limited by not checking the exact nature of shared data when it had to be requested from the authors or a repository. In some cases, reasonable requests might not be granted or the data shared may not include complete GWAS summary statistics. Data sharing outside of article texts, such as on project or author home pages, was also not checked.

## Conclusions

Only 10.6% of primary human GWAS studies either share or offer to share their summary statistics data in any form, which is low given that genomics is in many ways the leader in data sharing and this type of data is standardised, singled out for a NIH sharing mandate (we did not check whether the articles assessed in this study complied with funder mandates), has had recognised sharing value for over a decade, has public archives to host it, and often needs to be shared internally within research consortia. Other than potential human subjects ethics permissions issues, this type of data seems to be a best case for (partly) non-mandatory scientific data sharing. This suggests that data sharing is unlikely to become near-universal when it is optional. This emphasises the need for policy initiatives to promote data sharing, to extend the current apparently small minority of data sharing practices.

For GWAS as an illustration, formalised data sharing mandates implemented at the journal level would not be effective because GWAS studies can be published in general, health, psychology and medical journals in addition to specialist genetics and genomics journals. Alternative discipline-specific strategies may need to be devised, perhaps including agreements between funders for this type of data.

In terms of automatically identifying specific types of data reported to be shared in articles, the results suggest that in fields where data sharing statements are widely used, it should be possible to extract information about whether data was shared. Nevertheless, such sections seem to occur in less than 10% of articles in all broad fields of science (see data shared with: [49]) and so this strategy would not be widely effective. It is much more problematic to identify the type of data shared and seems impractical to automate this step. This is because data sharing statements are typically vague about what is shared and there is no single standard or policy adopted by all journals in a specific field regarding what should be included in a data access statement. Descriptions of the exact nature of the data available would not only help automation but also researchers scanning multiple articles to find relevant data for a new study. Thus, if more journals required data sharing statements and employed guidelines to ensure that the shared data was described in detail, or provided virtual rewards [50] for these activities, then this would support the level of automated data discovery that would be necessary to monitor data sharing and systematically identify shared data for later re-use.

## References

1. Krumholz, HM. Why data sharing should be the expected norm. BMJ, 350, h599.

2. Lindsay, DS. Sharing data and materials in Psychological Science. Psychological Science. 2017; 28(6): 699–702.

3. Mennes M, Biswal BB, Castellanos FX, Milham MP. Making data sharing work: the FCP/INDI experience. Neuroimage. 2013; 82: 683–691.

4. Wallis JC., Rolando E, Borgman CL. If we share data, will anyone use them? Data sharing and reuse in the long tail of science and technology. PloS ONE. 2013; 8(7): e67332. doi: 10.1371/journal.pone.0067332.

5. Burgess S, Scott RA, Timpson NJ, Smith GD, Thompson SG, EPIC-InterAct Consortium. Using published data in Mendelian randomization: a blueprint for efficient identification of causal risk factors. European Journal of Epidemiology. 2015; 30(7): 543–552.

6. Poline JB, Breeze JL, Ghosh SS, Gorgolewski K, Halchenko YO, Hanke M, et al. Data sharing in neuroimaging research. Frontiers in Neuroinformatics. 2012; 6, 9. doi: 10.3389/fninf.2012.00009.

7. Borgman CL. Big data, little data, no data: Scholarship in the networked world. Cambridge, MA: MIT Press; 2015.

8. Houtkoop BL, Chambers C, Macleod M, Bishop DV, Nichols TE, Wagenmakers EJ. Data sharing in psychology: A survey on barriers and preconditions. Advances in Methods and Practices in Psychological Science. 2018; 1(1): 70–85.

9. Nelson B. Data sharing: Empty archives. Nature News. 2009; 461(7261): 160–163.

10. Tenopir C, Allard S, Douglass K, Aydinoglu AU, Wu L, Read E, et al. (2011). Data sharing by scientists: practices and perceptions. PloS ONE. 2011; 6(6): e21101. doi: 10.1371/journal.pone.0021101.

11. Koslow SH. Sharing primary data: a threat or asset to discovery? Nature Reviews Neuroscience. 2002; 3(4): 311–313.

12. Tenopir C, Dalton ED, Allard S, Frame M, Pjesivac I, Birch B, et al. Changes in data sharing and data reuse practices and perceptions among scientists worldwide. PloS ONE. 2015; 10(8): e0134826. doi: 10.1371/journal.pone.0134826.

13. Mongeon P, Robinson-Garcia N, Jeng W, Costas R. Incorporating data sharing to the reward system of science: Linking DataCite records to authors in the Web of Science. Aslib Journal of Information Management. 2017; 69(5): 545–556.

14. Park H, You S, & Wolfram D. Informal data citation for data sharing and reuse is more common than formal data citation in biomedical fields. Journal of the Association for Information Science and Technology. 2018; 69(11): 1346–1354.

15. Peters I, Kraker P, Lex E, Gumpenberger C, Gorraiz J. Research data explored: an extended analysis of citations and altmetrics. Scientometrics. 2016; 107(2): 723–744.

16. Piwowar HA, Day RS, Fridsma DB. Sharing detailed research data is associated with increased citation rate. PloS ONE. 2007; 2(3): e308. doi: 10.1371/journal.pone.0000308.

17. Robinson-García N, Jiménez-Contreras E, Torres-Salinas D. Analyzing data citation practices using the data citation index. Journal of the Association for Information Science and Technology. 2016; 67(12): 2964–2975.

18. Stuart D. Data bibliometrics: metrics before norms. Online Information Review. 2017; 41(3): 428–435.

19. Sayogo DS, Pardo TA. Exploring the determinants of scientific data sharing: Understanding the motivation to publish research data. Government Information Quarterly. 2013; 30: S19–S31.

20. Fecher B, Friesike S, Hebing M. What drives academic data sharing? PloS ONE. 2015; 10(2): e0118053. doi: 10.1371/journal.pone.0118053.

21. Akers KG, Doty J. Disciplinary differences in faculty research data management practices and perspectives. International Journal of Digital Curation. 2013; 8(2): 5–26.

22. Demir E, Cary MP, Paley S, Fukuda K, Lemer C, Vastrik I, et al. The BioPAX community standard for pathway data sharing. Nature Biotechnology. 2010; 28(9): 935.

23. Thelwall M, Kousha K. Do journal data sharing mandates work? Life sciences evidence from Dryad. Aslib Journal of Information Management. 2017; 69(1): 36–45.

24. He L, Han Z. Do usage counts of scientific data make sense? An investigation of the Dryad repository. Library Hi Tech. 2017; 35(2): 332–342.

25. Zhao M, Yan E, Li K. Data set mentions and citations: A content analysis of full-text publications. Journal of the Association for Information Science and Technology. 2018; 69(1): 32–46.

26. Roche DG, Kruuk LE, Lanfear R, Binning, SA. Public data archiving in ecology and evolution: how well are we doing? PLoS Biology. 2015; 13(11): e1002295. doi: 10.1371/journal.pbio.1002295.

27. Khan N, Thelwall M, Kousha K. Does data sharing influence data reuse in biodiversity? A citation analysis. 23rd Nordic Workshop on Bibliometrics and Research Policy. 2018. Available from: https://figshare.com/articles/Does_Data_Sharing_Influence_Data_Reuse_in_Biodiversity_A_Citation_Analysis/7415312.

28. Doel T, Shakir DI, Pratt R, Aertsen M, Moggridge J, Bellon E, et al. GIFT-Cloud: A data sharing and collaboration platform for medical imaging research. Computer Methods and Programs in Biomedicine, 2017; 139(1): 181–190.

29. Groom Q, Weatherdon L, Geijzendorffer, IR. Is citizen science an open science in the case of biodiversity observations? Journal of Applied Ecology. 2017: 54(2): 612–617.

30. Costello MJ, Michener WK, Gahegan M, Zhang ZQ, Bourne, PE. Biodiversity data should be published, cited, and peer reviewed. Trends in Ecology & Evolution. 2013: 28(8), 454–461.

31. Beck J, Böller M, Erhardt A, Schwanghart W. Spatial bias in the GBIF database and its effect on modeling species’ geographic distributions. Ecological Informatics, 2014; 19: 10–15.

32. Ferguson AR, Nielson JL, Cragin MH, Bandrowski AE, Martone ME. Big data from small data: data-sharing in the ‘long tail’ of neuroscience. Nature neuroscience. 2014; 17(11): 1442–1447.

33. Boeckhout M, Zielhuis, GA, Bredenoord AL. The FAIR guiding principles for data stewardship: fair enough? European Journal of Human Genetics. 2018; 26(7): 931–936.

34. Guttmacher AE, Nabel EG, Collins FS. Why data-sharing policies matter. PNAS. 2009; 106(40): 16894.

35. Womack RP. Research data in core journals in biology, chemistry, mathematics, and physics. PloS ONE. 2015; 10(12): e0143460. doi: 10.1371/journal.pone.0143460.

36. Kaye J, Heeney C, Hawkins N, De Vries, Boddington. Data sharing in genomics—reshaping scientific practice. Nature Reviews Genetics. 2009; 10(5): 331–335.

37. Choudhury S, Fishman JR, McGowan ML, Juengst ET. Big data, open science and the brain: lessons learned from genomics. Frontiers in Human Neuroscience. 2014; 8: 239. doi: 10.3389/fnhum.2014.00239.

38. Birney E, Hudson, TJ, Green ED, Gunter C, Eddy S, Rogers J, et al. Prepublication data sharing. Nature. 2009; 461(7261): 168–170.

39. Shabani M, Borry P. “You want the right amount of oversight”: interviews with data access committee members and experts on genomic data access. Genetics in Medicine. 2016; 18(9): 892.

40. Shabani M, Dyke, SO, Joly Y, Borry P. Controlled access under review: improving the governance of genomic data access. PloS Biology. 2015; 13(12): e1002339. doi: 10.1371/journal.pbio.1002339.

41. Begum F, Ghosh D, Tseng GC, Feingold E. Comprehensive literature review and statistical considerations for GWAS meta-analysis. Nucleic acids research. 2012; 40(9): 3777–3784.

42. Bulik-Sullivan B, Finucane HK, Anttila V, Gusev A, Day FR, Loh PR, Daly MJ. An atlas of genetic correlations across human diseases and traits. Nature Genetics, 2015; 47(11): 1236–1241.

43. Evangelou E, Ioannidis JP. Meta-analysis methods for genome-wide association studies and beyond. Nature Reviews Genetics. 2013; 14(6): 379–389.

44. Bertagnolli MM, Sartor O, Chabner BA, Rothenberg ML, Khozin S, Hugh-Jones C, et al. Advantages of a truly open-access data-sharing model. 2017; NEJM, 376(12): 1178–1181.

45. NIH. Policy for Sharing of Data Obtained in NIH Supported or Conducted Genome-Wide Association Studies (GWAS). Available from: https://grants.nih.gov/grants/guide/notice-files/NOT-OD-07-088.html.

46. Buniello A, MacArthur JAL, Cerezo M, Harris LW, Hayhurst J, Malangone C, et al. The NHGRI-EBI GWAS Catalog of published genome-wide association studies, targeted arrays and summary statistics 2019. Nucleic Acids Research. 2019; 47 (Database issue): D1005–D1012. doi: 10.1093/nar/gky1120.

47. Goodman A, Pepe A, Blocker AW, Borgman CL, Cranmer K, Crosas M, et al. Ten simple rules for the care and feeding of scientific data. PloS Computational Biology. 2014; 10(4): e1003542. doi: 10.1371/journal.pcbi.1003542.

48. Savage CJ, Vickers, AJ. Empirical study of data sharing by authors publishing in PLoS journals. PloS ONE. 2009; 4(9): e7078. doi: 10.1371/journal.pone.0007078.

49. Thelwall M. The rhetorical structure of science? A multidisciplinary analysis of article headings. Journal of Informetrics. 2019; 13(3): 555–563.

50. Kidwell MC, Lazarević LB, Baranski E, Hardwicke TE, Piechowski S, Falkenberg LS, et al. Badges to acknowledge open practices: A simple, low-cost, effective method for increasing transparency. PloS Biology. 2016; 14(5): e1002456.

